# Adsorption mechanism of bacteria onto a Na-montmorillonite surface with organic and inorganic calcium

**DOI:** 10.1101/2020.10.08.332536

**Authors:** Guowang Tang, Cangqin Jia, Guihe Wang, Peizhi Yu, Xihao Jiang

## Abstract

The adsorption of bacteria onto the Na-montmorillonite (Na-MMT) was studied as a function of time, bacterial concentration, temperature and pH with the introduction of the organic and inorganic calcium sources. The results indicated that albeit revealing the same adsorption mechanism, the organic calcium (i.e., Ca(CH_3_COO)_2_) proposed in this study is more beneficial and environmentally friendly than the inorganic calcium (i.e., CaCl_2_) in terms of the adsorption of bacteria onto the Na-MMT surface, which can be ascribed to the formation of the denser aggregates in the Na-MMT with Ca(CH_3_COO)_2_. Meanwhile, the adsorption kinetics and isotherms followed the pseudo-second-order kinetic model and Langmuir Equation for both two calcium sources. Meanwhile, the adsorption bands of the water molecules on the minerals were observed to shift significantly after the bacterial adsorption, showing that the hydrogen bonding on the Na-MMT surface played an important role during this process. A value of ΔH^0^ > 0 indicated that the bacterial adsorption was affected by van der Waals force and hydrophobic interaction. Finally, the negative zeta potentials of the Na-MMT increased with the addition of Ca^2+^ ions, and the experimental data also showed that the adsorption of bacteria onto the Na-MMT was mainly determined by the electrostatic and non-electrostatic forces.

---

Compared with previous studies, Ca(CH_3_COO)_2_ was proposed for bacterial adsorption and adsorption mechanism of bacteria was clarified in the presence of Ca^2+^. Denser aggregates formed in the Ca(CH_3_COO)_2_ group explained its better adsorption capacity. Meanwhile, compared with CaCl_2_, Ca(CH_3_COO)_2_ was more environmentally friendly and eliminated the secondary pollution of Cl^-^ ions. Further, a new method to remove the bacteria from the aqueous solution was found.

## 1. Introduction

Bacterial pollution in drinking water is the main pollution. Contamination of bacteria can lead to food poisoning and disease. More than 1 million people die every year from water related diseases. At present, the size-exclusion mechanism is used as mainly way to remove contaminated bacteria in water sterilization and purification (1–4). The pore sizes of the cartilage or membranes need to be comparable to the bacterial size to remove bacteria in water. Despite all this, the membranes are easily clogged. Therefore, efficient methods and novel materials for water purification are highly needed. The complex material made of silver nanofibers, carbon nanotubes, and cotton have been widely studied as efficiency and fast purification speed (5–8). In this paper, we report a new principle demonstration of bacteria removal using Na-MMT based on a novel adsorption mechanism.

*Sporosarcina pasteurii*, which is a type of gram-negative and aerobic bacteria, has widely been applied in soil improvement by microbially induced calcium carbonate precipitation (MICP) (9–13), and some specific applications were found in practice such as the strengthening of soft-coal reservoirs (14), the improvement of the liquefaction resistance of sands (15, 16), the prevention and control of sandstorm (17–19) and the improvement of the environment as a dust suppressant (20). However, the porosity of soil could be blocked in the presence of bacteria, indicating the importance of the uniform distribution of bacteria within soil, which also leads to the extensive investigations of the adsorption of bacteria onto the mineral surface.

As aforementioned, numerous studies have focused on the adsorption of bacteria onto the mineral surface of the soil with various components such as Montmorillonite and Kaolinite (21, 22) as well as organic and inorganic mineral (23). Previous literature reported that the adsorption of bacteria onto the mineral surface played an important role in the bio-mineralization, bacterial distribution, bacterial activity and biodegradation (24). Meanwhile, the transport of bacteria controlled the migration of various pollutants including inorganic and organic pollutants as well as heavy metals. Thus, it is of great significance to understand the adsorption mechanism of bacteria onto the mineral surface, and several factors were identified to contribute to the sorption of bacteria (e.g., salt concentrations, pH, ionic strength and temperature) (24, 25). For instance, it is reported that also the adsorption capacity of bacteria onto the corundum was weakened with the growth of ionic strength and pH (23). Meanwhile, A. L. Mills et al. (26) found that the high salt concentrations led to more adsorption of bacteria onto the quartz surface, whereas D. Jiang et al. (27) demonstrated that the best attraction between bacteria and clay occurred when the temperature ranged from 15 to 35 °C.

Several studies confirmed that the calcium source is of great importance. CaCl_2_ was widely investigated and mainly used to change the ionic strength during the adsorption process (28–31). However, other calcium sources are also investigated by the researchers. J. Xu et al. (32) discovered that the biochemical properties of bacteria were strongly influenced by different calcium sources, and higher bacterial activity was found when the calcium lactate introduced in the MICP process. Meanwhile, the result also showed that the size and morphology of crystal were different in the presence of different calcium sources. Besides, the main component of the CaCO_3_ precipitates was identified as calcite when different calcium sources were used, and the results were also in line with another study (28). P. Li and W. Qu (31) demonstrated that the calcium acetate and calcium chloride presented the same effect on the repairing of the cracks and the strength boost of concrete in the MICP process, however, Y Zhang et al. (33) reported that the higher compressive strength of samples was obtained with the addition of calcium acetate source in porous media. Similarly, K. V Tittelboom et al. (34) used calcium acetate, calcium chloride and calcium nitrate to repair the cracked concrete, and the results presented that these three calcium sources revealed almost the same performance in view of the water permeability reduction.

It is well recognized that the durability of reinforced concrete was affected by the formation of cracks in concrete structures, which can be attributed to the degassed and electrochemistry corrosion caused by the chloride ions penetration through the cracks (35). Therefore, it is necessary to apply other organic calcium sources (e.g., Ca(CH_3_COO)_2_) instead of CaCl_2_, which will reduce the adverse effects owing to the Cl^-^ ions penetration within the samples, albeit CaCl_2_ has been widely studied for the adsorption of bacteria on the minerals. However, the effect of different calcium sources was rarely studied, and the detailed processes in terms of the adsorption of bacteria onto the mineral surface involving the Ca^2+^ ions was still unknown.

Given the foregoing, with different calcium sources introduced, the adsorption of bacteria onto the Na-montmorillonite (Na-MMT) surface was studied with several parameters investigated including the time, bacterial concentration, temperature and pH by batch experiments. At the same time, the desorption of bacteria on the Na-MMT was also investigated. Further, the adsorption mechanism of bacteria onto the Na-MMT surface containing different calcium sources was clarified by means of Brunner–Emmet–Teller (BET), Fourier Transform infrared spectroscopy (FTIR) and the Zeta potential measurement. Finally, the morphology of bacteria adsorbed on the Na-MMT surface was also facilitated by the use of Cryo-SEM.

## 2. Materials, preparation of bacteria and Methodologies

### 2.1 Materials

#### 2.1.1 *Bacterium* and Culturing

The bacteria used was purchased from Shanghai Fusheng industrial Reagent Co., Ltd, and was characterized as a strain of *Sporosarcina pasteurii*. The colonies of the bacteria were inoculated into 1000 mL of CASO+20 g/L urea medium and were shaken (120 rev minutes^−1^) at 301.15 K for 24 hours. The culture medium was centrifuged at 8000 rev minutes ^−1^ at 277.15 K for 10 min to obtain the bacteria. Then, the bacteria were washed for three times by the sterilized distilled-deionized (DDI) water and were resuspended in DDI. Then 8.101 mol L^−1^ CaCl_2_ and Ca(CH_3_COO)_2_ solution was prepared to obtain a known concentration of bacterial suspension. The concentration of Ca^2+^ ion was optimized through experiments. The optical density (OD) of bacteria at 600 nm wavelength was analyzed by a UV-vis spectrophotometer (UV-752, China), and the bacterial concentration followed an optical density at 600 nm (OD_600_).

#### 2.1.2 Mineral

The Na-MMT was purchased from Zhejiang Fenghong new materials Co., Ltd. The physical and chemical properties of the Na-MMT were characterized by XRD, FTIR and XRF, as listed in Fig. S1 (A), Fig. S1 (B), Table S1and Table S2, respectively.

### 2.2 Preparation of Bacteria in the Adsorption and Desorption Experiments

A series of sorption experiments was performed to investigate the adsorption of bacteria on the Na-MMT surface with CaCl_2_ and Ca(CH_3_COO)_2_ introduced as a function of time, bacterial concentration, temperature and pH. The bacteria were centrifuged and resuspended (OD_600_=1.5) by 8.101 mol L^−1^ CaCl_2_ and Ca(CH_3_COO)_2_ solution, respectively. The mixture of 0.4 grams of Na-MMT and 100 grams of suspension solution (OD_600_=1.5) was stirred at 240 rev minutes^−1^ for 20 min to investigate the effect of the time change (i.e., 0 min to 120 min), temperature (i.e., 293 K - 333 K) and pH (i.e., 7-11) on the bacterial adsorption onto the mineral. To study the influence of the initial bacterial concentrations on the adsorption of the bacteria, the mixture of 0.4 grams of Na-MMT and 100 grams of suspension solution (OD_600_=0.5, 1.0, 1.5 and 2.0) was stirred at 240 rev minutes^−1^ for 20 minutes. The amount of bacterial adsorbed onto the mineral was calculated by subtracting the current bacterial concentration from the initial amount of bacteria added (without any bacterial adsorption).

The mixture of 0.4 grams of Na-MMT and 100 grams of suspension solution prepared by DDI (OD600=0.5, 1.0 and 1.5) was stirred at 240 rev minutes^−1^ for 20 minutes. The concentration of bacteria (D1) was measured until the values were stable. The desorption experiments of bacteria were conducted by shaking the suspension solution at 120 rev minutes^−1^ for 2 hours. The concentration of bacteria (D2) was then measured again until no significant changes occurred in the values. The percentage of bacterial desorption (W) was calculated by Eq. (1).

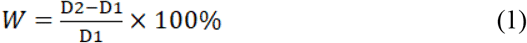

### 2.3 Methodologies

#### 2.3.1 Fourier Transform Infrared Spectra (FTIR)

FTIR (Thermo Scientific Nicolet iS5) was employed to characterize the adsorption mechanism of bacteria onto the Na-MMT surface. The mixture of 2 mg of powder sample and 200 mg of pure KBr was ground evenly and was then placed into the mold. The mixture was pressed into a transparent sheet by the hydraulic press and was put into the infrared spectrometer for the test with a wavenumber range of 4000-400cm^-1^, scanning times of 32 and a resolution of 4cm^-1^.

#### 2.3.2 Cryo-Scanning Electron Microscope (SEM)

Cryo-scanning electron microscope (Cryo-SEM, FEI Quanta 450) was used to observe the image of the Na-MMT surface with the introduction of calcium sources. The powder samples were frozen for 30 seconds in liquid nitrogen snow mud, and then was sputtered with 10 mA current for 60 seconds, after sublimation at 363 K for 10 minutes. Then, the platinum was sprayed on the surface of the sample. Finally, the sample was involved in SEM for observation with a 5 kV of accelerating voltage.

#### 2.3.3 Electrokinetic and Surface Characterization of Mineral

The minerals (i.e., Na-MMT, Na-MMT with Ca(CH_3_COO)_2_ and CaCl_2_) were diluted in the deionized water to ensure the final concentration between 5-10 mg mL^−1^. Then, the Zeta-potential values of minerals were measured at 298 K by a zeta potential analyzer (Malvern Zetasizer Nano ZS90). All experiments were repeated for 3 times at a pH of 8. The specific surface area and the adsorption cumulative volume of pores within the minerals were obtained by N_2_ adsorption (Mike 2020).

## 3. Results

### 3.1 Adsorption and Desorption of Bacteria onto the Na-MMT Surface

The amount of bacterial adsorption onto the Na-MMT surface at a pH of 8.5 was almost the same when different calcium sources (i.e., CaCl_2_ and Ca(CH_3_COO)_2_) were introduced (Fig. 1 (a)). The adsorption of bacteria proceeded rapidly in the first 30 minutes but the process slowed down after 30 minutes. Eventually, the bacteria were completely adsorbed onto the Na-MMT surface. It may be due to a change in the adsorption process between the surfaces with bacteria and mineral. Because the adsorption of bacteria is driven by the release of counterions to the charges on the bacteria and the surface (36)

**Figure 1.**
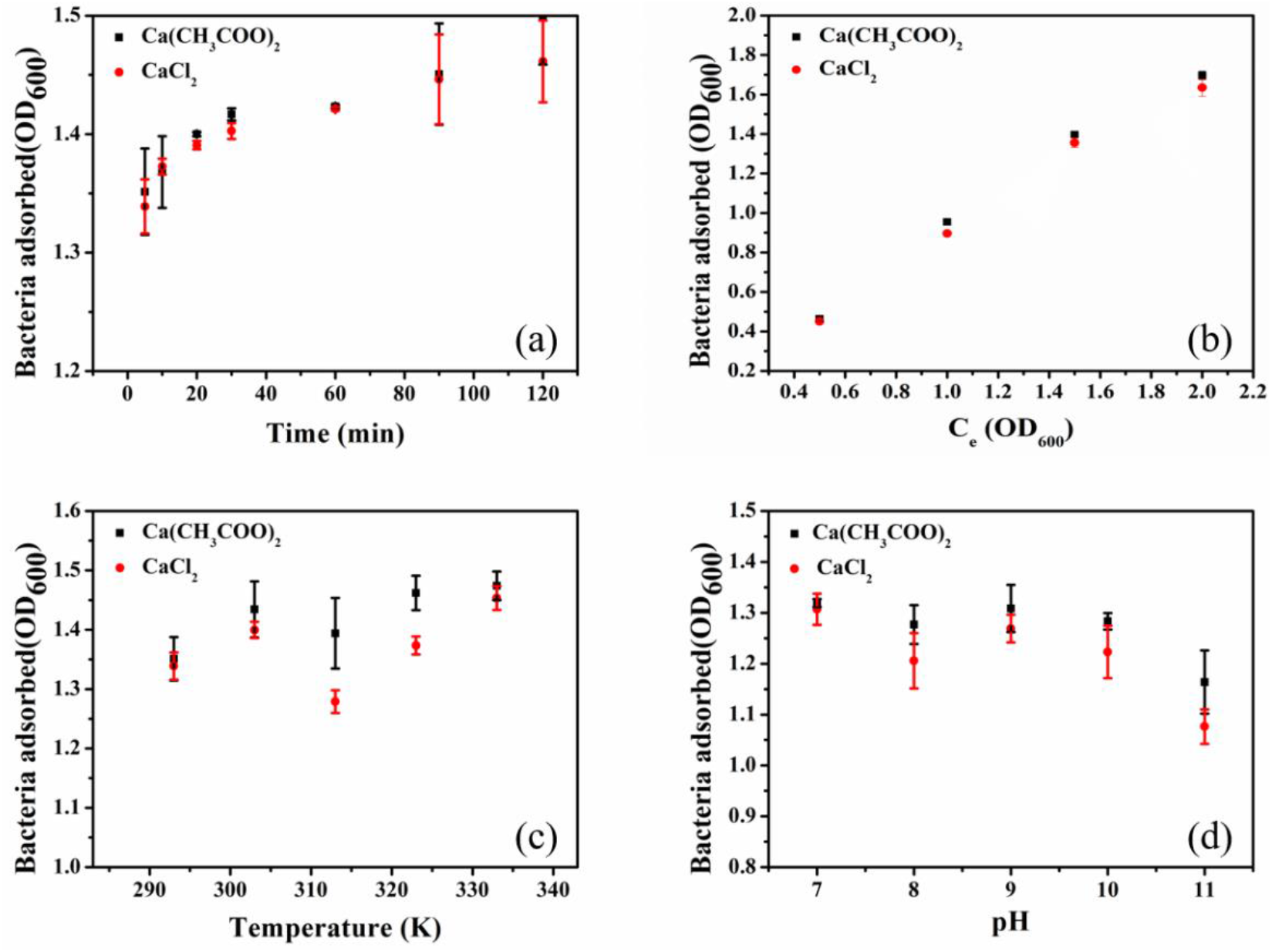
Effect of the contact time (a), bacterial concentration (b), temperature (c) and pH (d) on the adsorption of bacteria onto a Na-MMT surface with 8.1 mmol/L of Ca^2+^

The amount of bacterial sorption on the Na-MMT surface increased with the growth of bacteria concentration (Fig. 1 (b)). When the OD_600_ of bacteria were 0.5, 1.0 and 1.5, the bacteria were almost completely absorbed under different calcium sources, whereas when the OD_600_ of bacteria was 2.0, the percentage of bacteria adsorbed by Na-MMT was only 86.5 % and 83.5 % with the introduction of Ca(CH_3_COO)_2_ and CaCl_2_, respectively. The reason could be that adsorption capacity of bacteria onto Na-MMT surface has reached the saturation value.

Fig. 1 (c) shows that the adsorption of bacteria on the Na-MMT surface with different calcium sources were dependent on the temperature. A significant change was observed with the increment of the temperature. However, instead of an increase, the number of bacterial sorption decreased at 313 K, and further investigations should be performed with respect to this trend.

The adsorption of bacteria on the Na-MMT surface was significantly affected by the pH under different calcium sources, as shown in Fig. 1(d)). With CaCl_2_ and Ca(CH_3_COO)_2_ introduced, when the pH increased from 7 to 11, a gradual reduction in the bacterial adsorption onto the Na-MMT surface was seen. Meanwhile, compared to CaCl_2_, a larger volume of bacteria were adsorbed onto the mineral surface with the introduction of Ca(CH_3_COO)_2_, as shown in Fig. 1, which could be attributed to the fact that the growth of pH led to the increase of electrostatic repulsion both on the surface of minerals and the bacteria (24).

The mixture of bacteria and Na-MMT was shaken at 120 rpm for 2 hours to determine the percentage of sorption for bacteria. Then, the percentage of desorption for bacteria has been calculated. When the OD_600_ was 0.5, 1.0 and 1.5, nearly no bacteria were released from the Na-MMT surface with different calcium sources introduced (the data was shown in Fig. 2S), thus suggesting that the bacteria were strongly retained by the Na-MMT surface in the presence of Ca^2+^ ions. This phenomenon is attributed to strong adsorption of bacteria on mineral by chemical interaction.

**Figure 2.**
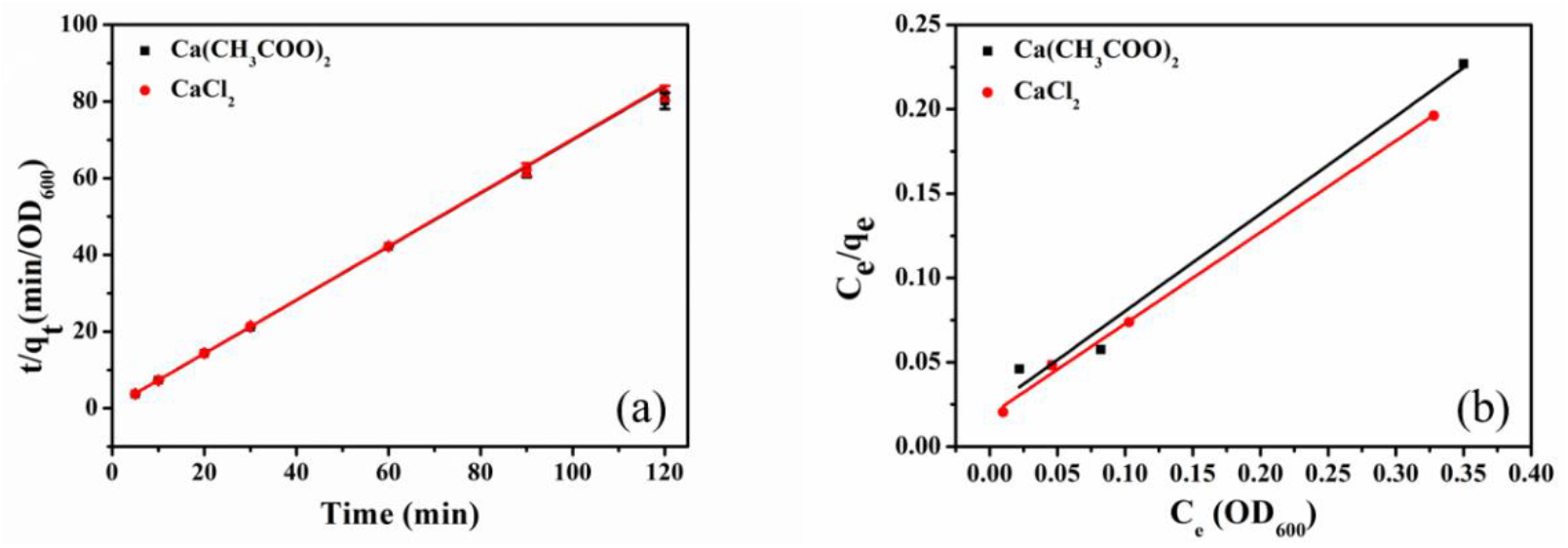
Pseudo-second-order kinetic models (a) and Langmuir isotherm models (b) for the adsorption of the bacteria onto the Na-MMT surface with 8.1 mmol/L of Ca^2+^

### 3.2 Kinetic and Isotherm of Bacteria Adsorption

The pseudo-second-order kinetic model was applied in terms of the adsorption of bacteria onto the Na-MMT surface with different calcium sources introduced (Fig. 2(a)). The experimental data were in line with a pseudo-second-order kinetic model (Table. S3). The correlation coefficients (R^2^ > 0.99) indicate that the sorption process of bacteria onto the Na-MMT surface may be ascribed to chemisorption (37, 38). Meanwhile, the results also suggest that when different calcium sources were introduced, the overall rate of the adsorption process was controlled by the chemical adsorption when the bacteria were present (39, 40).

With different calcium sources introduced, the adsorption of bacteria onto the Na-MMT surface followed a Langmuir isotherm model (Fig. 2(b)). The experiment data fitted well with the Langmuir isotherm model (Table. S3). In the Na-MMT-Ca(CH_3_COO)_2_ group, the Langmuir constant K_L_ was 12.2 % smaller than that of Na-MMT-CaCl_2_ group, indicating that more energy was released with respect to the adsorption of bacteria where Ca(CH_3_COO)_2_ was added, in comparison with CaCl_2_. In view of the above findings, it can be inferred that the sorption capacity of the Na-MMT involving Ca(CH_3_COO)_2_ was greater than that with CaCl_2_. The detailed explanation will be presented in section result.

### 3.3 FTIR Analysis of Adsorption

The FTIR analysis of selected samples was exhibited in Fig. 3. The adsorption bands of the mineral were analyzed as follows. The peak 1040 cm^−1^ in the curves can be assigned to the Si-O stretching vibrations, whereas the peaks, 469, 523 and 917 cm^−1^ were due to the Si-O-Si bending vibration. For the bacteria, CH_2_ asymmetric stretching vibration was observed at 2936 cm^-1^, and the presence of C=O contributed to the peak 1655 cm^-1^. The peak 1403 cm^-1^ and 1059 cm^-1^ can be ascribed to the presence of C-O bending and polysaccharide. The experimental data indicated that the Si-O stretching vibrations and the Si-O-Si bending vibration of the Na-MMT were almost the same with the introduction of CaCl_2_ or Ca(CH_3_COO)_2_. Besides, the peaks 3410 cm^-1^ (3449 cm^-1^) and 1641 cm^-1^ (1642 cm^-1^) were due to the symmetric stretching and bending vibrations of the water molecules on the Na-MMT-Ca(CH_3_COO)_2_ group (Na-MMT-CaCl_2_ group). Meanwhile, the FTIR curve also shows that after the sorption of bacteria, the functional group position of the water molecules on the Na-MMT-Ca(CH_3_COO)_2_ group (Na-MMT-CaCl_2_ group) shifted from 3420 to 3424 cm^-1^ (3440 to 3422 cm^-1^) and from 1641 to 1645 cm^-1^ (1642 to 1654 cm^-1^), respectively. In view of the aforementioned findings, it is suggested that the water molecules on the Na-MMT surface played an important role in the sorption process. Moreover, the CH_2_ asymmetric stretching vibration and the C=O were observed on the mineral-Ca(CH_3_COO)_2_-bacteria and mineral-CaCl_2_-bacteria, confirming the adsorption of bacteria onto the Na-MMT surface, and Ca(CH_3_COO)_2_ and CaCl_2_ show a similar mechanism with regard to the adsorption of bacteria onto the Na-MMT surface. The wavelength number of mineral and bacteria was shown in Table. S4.

**Figure 3.**
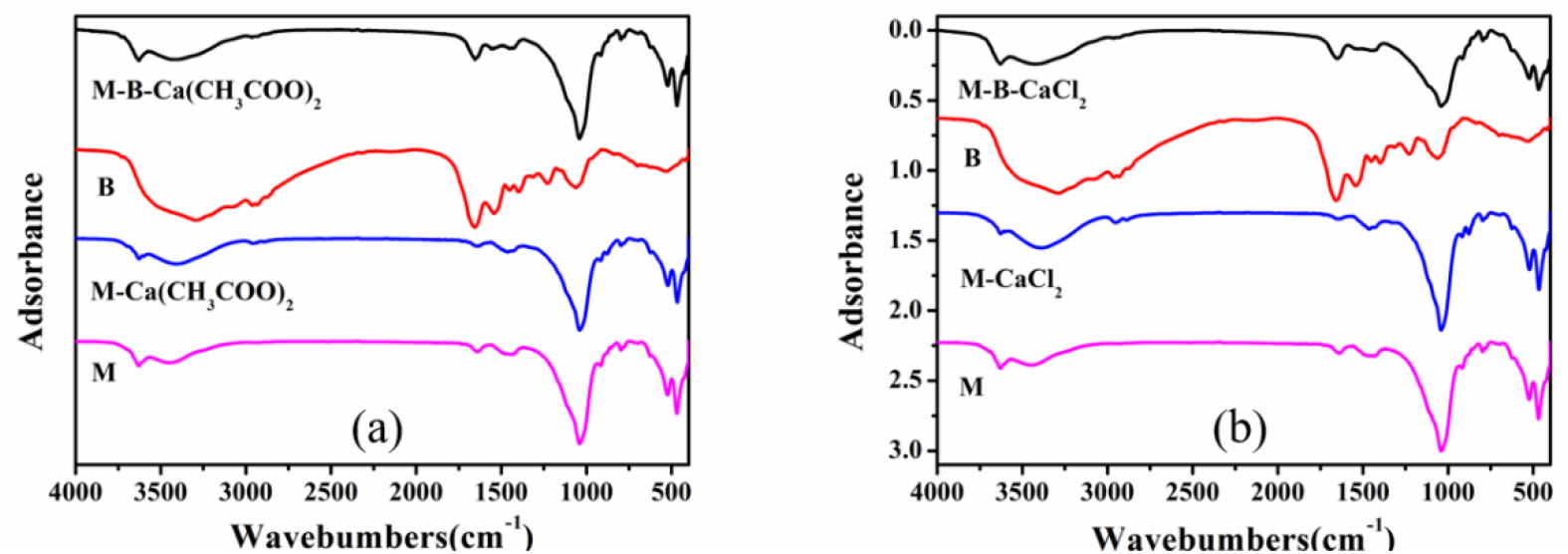
FTIR spectra of Na-MMT, Na-MMT-Ca(CH_3_COO)_2_, Na-MMT-CaCl_2_, bacteria, Na-MMT-Ca(CH_3_COO)_2_-bacteria and Na-MMT-CaCl_2_-bacteria (M, Na-MMT; B, bacteria)

### 3.4 SEM Analysis of Bacterial Adsorption

The images of bacterial adsorption on the Na-MMT surface were observed by the use of Cryo-SEM (Fig. 4). Compared to CaCl_2_ (Fig. 4(c) and 4(d)), the images suggested that Ca(CH_3_COO)_2_ presented a tendency to aggregate the Na-MMT and led to the formation of larger particles during the bacterial adsorption process (Fig. 4(a) and 4(b)). The bacteria were mainly attached onto the Na-MMT surface in the presence of Ca(CH_3_COO)_2_ and CaCl_2_. The images (Fig. 4(b) and (c)) indicate that although different calcium sources were introduced, the adsorption mechanism of bacteria onto the Na-MMT surface was similar.

**Figure 4.**
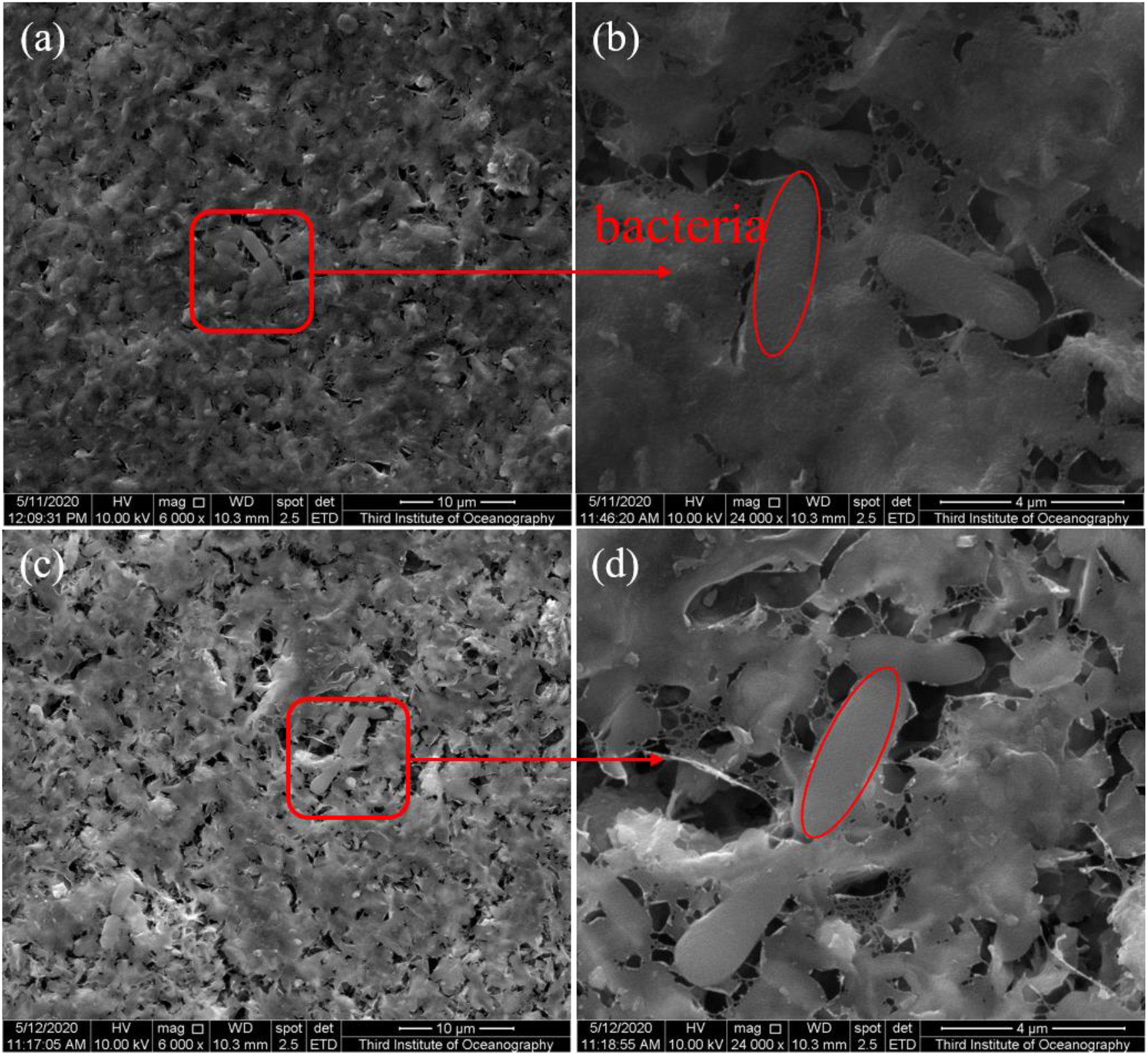
Images of the bacterial adsorption onto the Na-MMT surface with the addition of Ca(CH_3_COO)_2_ (a and b) and CaCl_2_ (c and d)

### 3.5 BET Analysis of Bacterial Adsorption

The specific surface area of the Na-MMT increased in the presence of CaCl_2_ and Ca(CH_3_COO)_2_ (Fig. 5(a)). The results showed that the specific surface area affects the adsorption of bacteria onto the Na-MMT surface to some degree. The Na-MMT with Ca(CH_3_COO)_2_ adsorbed a greater amount of bacteria than that with CaCl_2_, although the Na-MMT with Ca(CH_3_COO)_2_ presents a smaller SSA than that with CaCl_2_. Therefore, considering this, it is implied that the specific surface area may not be the major contributor to the bacterial adsorption onto the Na-MMT surface, which could be related to the non-electrostatic force dominating the bacteria adsorption onto the Na-MMT surface. Meanwhile, the hysteresis loop of the isotherm adsorption line of mineral followed H3 type. Besides, the results also indicated that the adsorption characteristics of the Na-MMT failed to change significantly with different calcium sources introduced (i.e., CaCl_2_ and Ca(CH_3_COO)_2_). Further, it is demonstrated that a large number of pores were seen in these minerals (Fig. 5(b)), and the cumulative volume of the pores within the Na-MMT increased with the introduction of Ca(CH_3_COO)_2_, while a reduction was seen in the pore volume within the mineral with CaCl_2_. Therefore, the cumulative volume of pores within the Na-MMT might not be a major factor as well in the determination of the bacteria adsorption onto the mineral.

**Figure 5.**
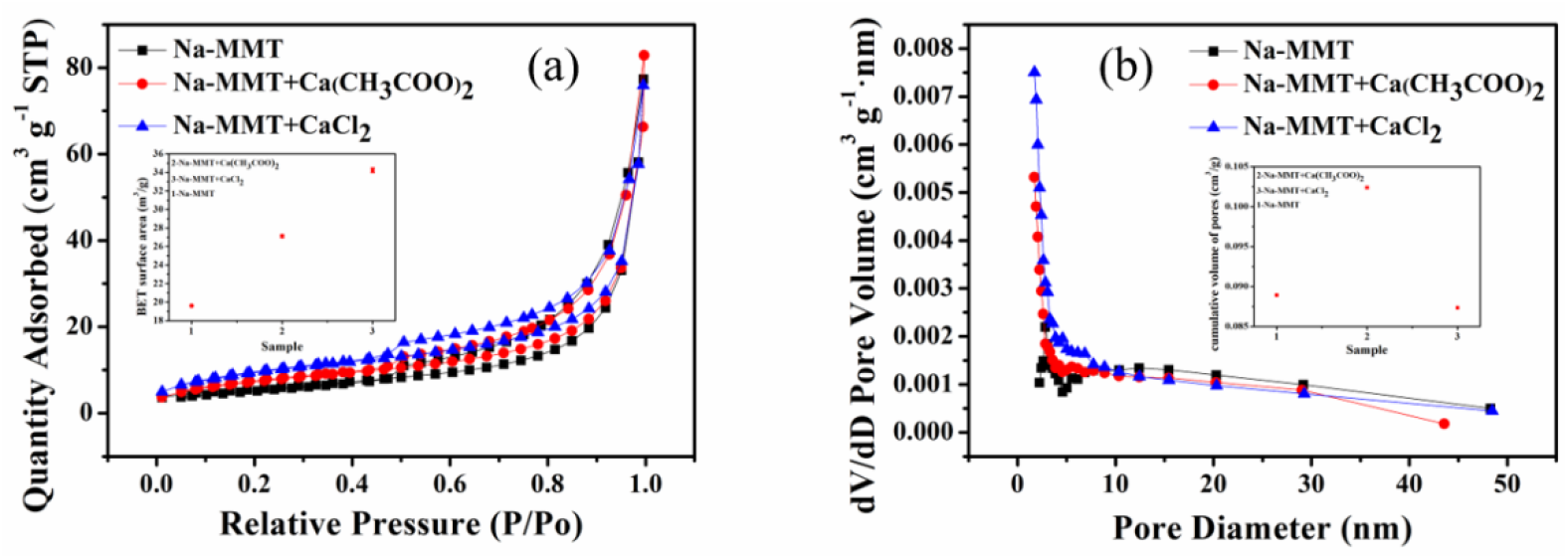
Specific surface area and cumulative volume of pores of minerals (Na-MMT, Na-Ca(CH_3_COO)_2_ and Na-MMT-CaCl_2_)

## 4. Discussions

Absorption refers to the process by which one material occupies another one through the small pores or spaces between them, which involves the whole volume of materials, whereas adsorption is defined as the process where the atoms, ions or molecules from a gas, liquid or dissolved solid adhere to the surface, and only the surface area of material is involved. Due to the small interlayer spacing of Na-MMT (i.e., 0-10 nm) (41, 42) and the larger size of bacteria (3-5 um), the bacteria were only thought to be adsorbed on the Na-MMT surface instead of throughout the whole mineral. Moreover, the experimental data show the amount of bacteria adsorbed on the Na-MMT surface increase with increasing temperature, which is a typical adsorption phenomenon. Langmuir Equation was well fitted by the experimental data, which confirms that the adsorption of bacteria onto the Na-MMT surface was single molecular layer adsorption. The results of FTIR spectra showed that the vibration peak of water molecules was significantly different during the adsorption process, proving that the hydrogen bonds played an important role.

The Fig 6 shows that the zeta potential shifted from −26.3 mv to −13 mv in the presence of Ca(CH_3_COO)_2_, whereas a similar shift was also observed in the presence of CaCl_2_ (i.e., from −26.3 mv to −14.2 mv), accounting for the volume increase of bacteria adsorbed onto the Na-MMT surface due to the introduction of Ca^2+^ ions. The absolute value of zeta potential on the Na-MMT surface in the presence of Ca(CH_3_COO)_2_ (−13 mv) was smaller than that CaCl_2_ (−14.2 mv), explaining the superior performance of Ca(CH_3_COO)_2_ than CaCl_2_ regarding the adsorption of bacteria onto the Na-MMT surface. The adsorption of bacteria on the minerals is mainly affected by the electrostatic and non-electrostatic forces. The electrostatic force is generated by the Coulomb force interaction between two charged substances, while the non-electrostatic force is generated by the Van der Waals force, Hydrophobic interaction and Hydrogen bond. Due to the negative charge of Na-MMT surface and bacteria, we considered that electrostatic force is not conducive to bacterial adsorption on the Na-MMT surface. The experimental data showed that the addition of Ca^2+^ ions significantly increased the number of bacteria adsorbed on the Na-MMT surface. With the introduction of Ca^2+^ ions, the negative charges of the Na-MMT surface decreased significantly (Fig. 6). Therefore, the above findings also imply that the non-electrostatic force may dominate in the bacterial adsorption onto the Na-MMT surface compared with the electrostatic interaction. K. S. Zerda et al. (43) and S. Chattopadhyay and R. W. Puls (44) also found that virus adsorption on silica and the adsorption of bacteriophages on and kaolinite were mainly govern by the non-electrostatic forces. The experimental data of FTIR showed an obvious change was observed in the functional groups of the water molecules on the mineral after the bacterial adsorption. As reported, the hydrogen bonds between two interacting bodies were formed by these shifts (45, 46).

**Figure 6.**
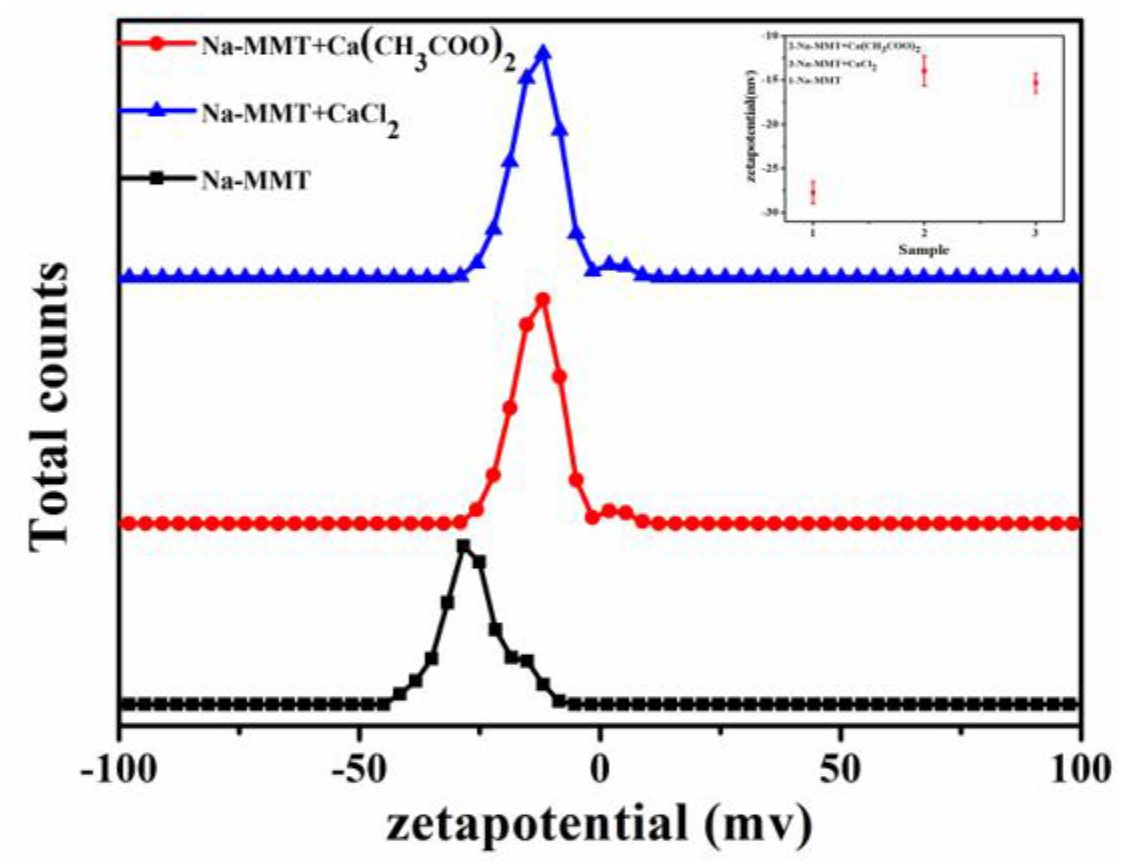
Zeta-potential values of minerals (Na-MMT, Na-MMT-Ca(CH_3_COO)_2_ and Na-MMT-CaCl_2_)

The adsorption of bacteria onto the mineral surface across a range of temperatures (303 to 333 K) with addition of 8.1 mmol/L Ca^2+^ ions is shown in Fig 7. The parameter values of ΔS^0^ and ΔH^0^ can be calculated through the slope and intercept, respectively. The thermodynamics parameter values of Na-MMT are listed in Table S5. As ΔG^0^ < 0, the adsorption of bacteria onto the Na-MMT surface was spontaneous. P. D. Ross and S. Subramanian (47) reported that the hydrogen bond may cause negative enthalpy, while ion interaction and hydrophobic interaction may result in positive enthalpy. A value of ΔH^0^ > 0 indicated that the bacterial adsorption was affected by van der Waals force and hydrophobic interaction. Thus, confirming that the non-electrostatic forces presented an important effect on the adsorption of bacteria onto the Na-MMT surface. The adsorption process of bacteria onto Na-MMT surface is schematically illustrated in Fig 8.

**Figure 7.**
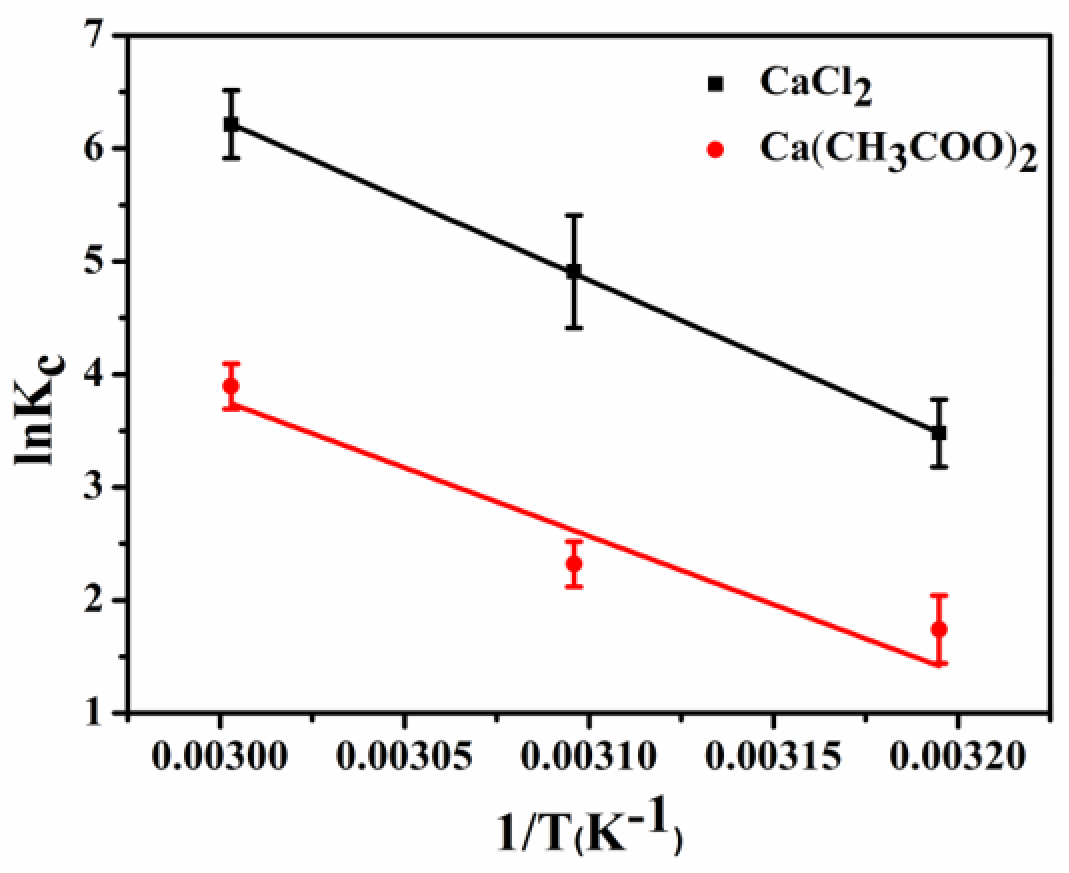
Thermodynamics of the adsorption of the bacteria on the Na-MMT surface with the addition of 8.1 mol L^-1^ Ca(CH_3_COO)_2_ and CaCl_2_.

**Figure 8.**
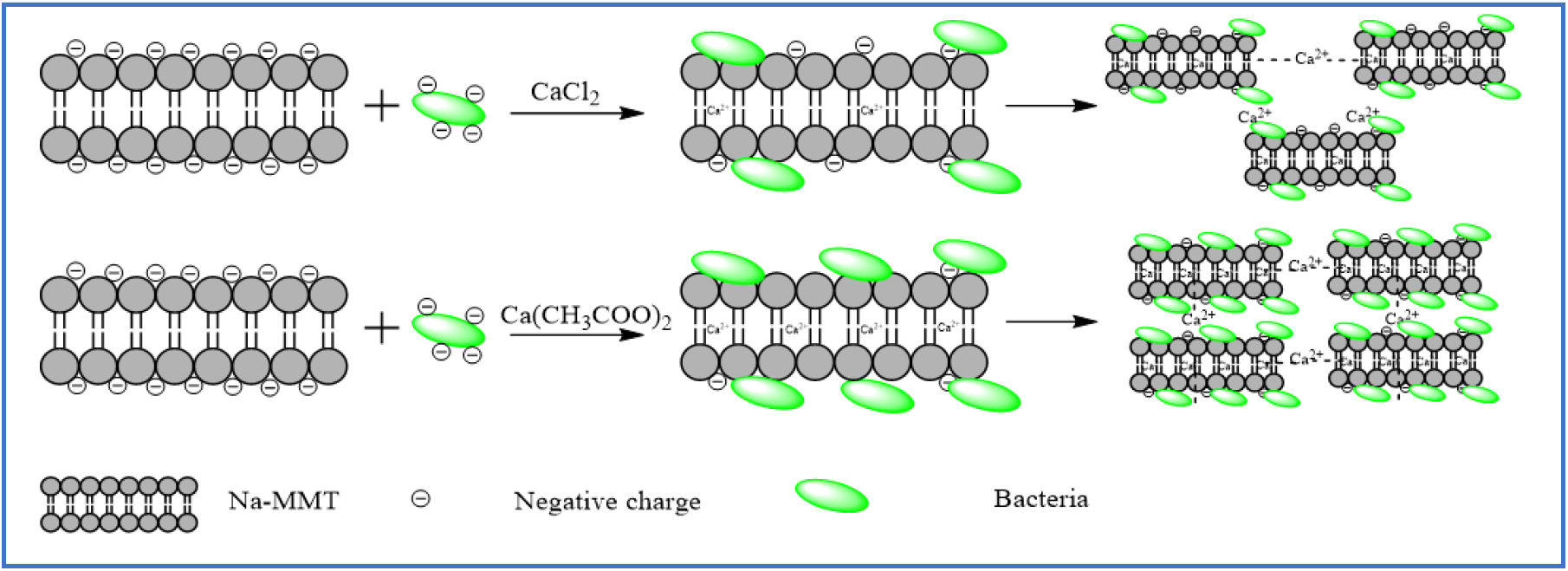
Schematic illustration of the bacterial adsorption process onto the Na-MMT in the presence of Ca^2+^.

The growth of pH increased the negative charge both on the surface of minerals and the bacteria (24). Due to the increment of the repulsive electrostatic force, a reduction was seen in the bacteria adsorption onto the mineral surface as pH increased. The images of SEM showed that the aggregates of the Na-MMT involving Ca(CH_3_COO)_2_ were much denser than that with CaCl_2_, indicating that in comparison with CaCl_2_, a larger volume of bacteria were adsorbed onto the mineral with Ca(CH_3_COO)_2_, which can be assigned to the smaller value of the Langmuir parameter K_L_ in this group, indicating a stronger affinity of bacteria for the Na-MMT containing Ca(CH_3_COO)_2_. Almost no bacteria were desorbed, which could be assigned to the chemical interaction that could result in strong adsorption of bacteria onto the mineral surfaces and low mobility of those adsorbed bacteria.

## 5. Conclusions

As the Na-MMT with Ca(CH_3_COO)_2_ presented a higher affinity for the bacteria than CaCl_2_, Ca(CH_3_COO)_2_ outperformed CaCl_2_ in terms of the bacterial adsorption onto the Na-MMT surface. The adsorption of bacteria onto the Na-MMT surface was identified as chemical adsorption, and it is mainly governed by the non-electrostatic forces (i.e., the van der Waals force, hydrophobic interaction and hydrogen bonding) and electrostatic forces with addition of Ca^2+^. The specific surface area of Na-MMT increased with the addition of Ca^2+^ ions, and the adsorption of bacteria was also affected by the specific surface area of minerals, but it is not a main contributor.

## Acknowledgement

This work was financially supported by Research Foundation of Key Laboratory of Deep Geodrilling Technology, Ministry of Natural Resources [No. KF2019X] and Fouling mechanism and control method of inner wall of core drill pipe [No. 41772388].

## References

1. Liu S, Zeng TH, Hofmann M, Burcombe E, Wei J, Jiang R, Kong J, Chen Y 2011. Antibacterial Activity of Graphite, Graphite Oxide, Graphene Oxide, and Reduced Graphene Oxide: Membrane and Oxidative Stress. Acs Nano 5:6971–6980.

2. Song WH, Ryu HS, Hong SH. 2008. Antibacterial properties of Ag (or Pt)-containing calcium phosphate coatings formed by micro-arc oxidation. Journal of Biomedical Materials Research Part A 88A:246–254.

3. Wieprecht T, Apostolov O, Beyermann M, Seelig J. 2000. Membrane Binding and Pore Formation of the Antibacterial Peptide PGLa: Thermodynamic and Mechanistic Aspects †. Biochemistry 39:442–452.

4. Zeng X, Mccarthy DT, Deletic A, Zhang X. 2015. Silver/Reduced Graphene Oxide Hydrogel as Novel Bactericidal Filter for Point-of-Use Water Disinfection. Advanced Functional Materials 25:4344–4351.

5. Yoon KY, Byeon JH, Park CW, Hwang J. 2008. Antimicrobial Effect of Silver Particles on Bacterial Contamination of Activated Carbon Fibers. Environmental Science & Technology 42:1251–1255.

6. Schoen DT, Schoen AP, Hu L, Kim HS, Heilshorn SC, Cui Y. 2010. High Speed Water Sterilization Using One-Dimensional Nanostructures. Nano Letters 10:3628–3632.

7. Birbir Y, Ur G, Birbir M. 2008. Inactivation of bacterial population in hide-soak liquors via direct electric current. Journal of Electrostatics 66:355–360.

8. Jain P, Pradeep T. 2005. Potential of silver nanoparticle-coated polyurethane foam as an antibacterial water filter. Biotechnology & Bioengineering 90:59–63.

9. Cheng L, Cord-Ruwisch R. 2014. Upscaling Effects of Soil Improvement by Microbially Induced Calcite Precipitation by Surface Percolation. Geomicrobiology Journal 31:396–406.

10. DeJong JT, Mortensen BM, Martinez BC, Nelson DC. 2010. Bio-mediated soil improvement. Ecological Engineering 36:197–210.

11. García-González J, Rodríguez-Robles D, Wang J, De Belie N, Morán-del Pozo JM, Guerra-Romero MI, Juan-Valdés A. 2017. Quality improvement of mixed and ceramic recycled aggregates by biodeposition of calcium carbonate. Construction and Building Materials 154:1015–1023.

12. Kakelar MM, Ebrahimi S, Hosseini M. 2016. Improvement in soil grouting by biocementation through injection method. Asia-Pacific Journal of Chemical Engineering 11:930–938.

13. Umar M, Kassim KA, Ping Chiet KT. 2016. Biological process of soil improvement in civil engineering: A review. Journal of Rock Mechanics and Geotechnical Engineering 8:767–774.

14. Song C, Elsworth D. 2018. Strengthening mylonitized soft-coal reservoirs by microbial mineralization. International Journal of Coal Geology 200:166–172.

15. Xiao P, Liu H, Xiao Y, Stuedlein AW, Evans TM. 2018. Liquefaction resistance of bio-cemented calcareous sand. Soil Dynamics and Earthquake Engineering 107:9–19.

16. Zamani A, Montoya BM. 2018. Undrained Monotonic Shear Response of MICP-Treated Silty Sands. Journal of Geotechnical and Geoenvironmental Engineering 144:04018029.

17. Maleki M, Ebrahimi S, Asadzadeh F, Emami Tabrizi M. 2016. Performance of microbial-induced carbonate precipitation on wind erosion control of sandy soil. International Journal of Environmental Science and Technology 13:937–944.

18. Tian K, Wu Y, Zhang H, Li D, Nie K, Zhang S. 2018. Increasing wind erosion resistance of aeolian sandy soil by microbially induced calcium carbonate precipitation. Land Degradation & Development 29:4271–4281.

19. Wang Z, Zhang N, Ding J, Lu C, Jin Y. 2018. Experimental Study on Wind Erosion Resistance and Strength of Sands Treated with Microbial-Induced Calcium Carbonate Precipitation. Advances in Materials Science and Engineering 2018:1–10.

20. Naeimi M, Chu J. 2017. Comparison of conventional and bio-treated methods as dust suppressants. Environ Sci Pollut Res Int 24:23341–23350.

21. Hong Z, Rong X, Cai P, Dai K, Liang W, Chen W, Huang Q. 2012. Initial adhesion of Bacillus subtilis on soil minerals as related to their surface properties. European Journal of Soil Science 63:457–466.

22. Zhou X, Huang Q, Chen S, Yu Z. 2005. Adsorption of the insecticidal protein of Bacillus thuringiensis on montmorillonite, kaolinite, silica, goethite and Red soil. Applied Clay Science 30:87–93.

23. Zhao WQ, Liu X, Huang QY, Rong XM, Liang W, Dai K, Cai P. 2012. Sorption of Streptococcus suison various soil particles from an Alfisol and effects on pathogen metabolic activity. European Journal of Soil Science 63:558–564.

24. Rong X, Huang Q, He X, Chen H, Cai P, Liang W. 2008. Interaction of Pseudomonas putida with kaolinite and montmorillonite: a combination study by equilibrium adsorption, ITC, SEM and FTIR. Colloids Surf B Biointerfaces 64:49–55.

25. Hong Z, Rong X, Cai P, Liang W, Huang Q. 2011. Effects of Temperature, pH and Salt Concentrations on the Adsorption ofBacillus subtilison Soil Clay Minerals Investigated by Microcalorimetry. Geomicrobiology Journal 28:686–691.

26. Mills AL, Herman JS, Hornberger GM, Dejesús TH. 1994. Effect of Solution Ionic Strength and Iron Coatings on Mineral Grains on the Sorption of Bacterial Cells to Quartz Sand. Applied & Environmental Microbiology 60:3300–3306.

27. Jiang D, Huang Q, Cai P, Rong X, Chen W. 2007. Adsorption of Pseudomonas putida on clay minerals and iron oxide. Colloids & Surfaces B Biointerfaces 54:217–221.

28. De Muynck W, Cox K, Belie ND, Verstraete W. 2008. Bacterial carbonate precipitation as an alternative surface treatment for concrete. Construction and Building Materials 22:875–885.

29. Pacheco-Torgal F, Labrincha JA. 2013. Biotech cementitious materials: Some aspects of an innovative approach for concrete with enhanced durability. Construction & Building Materials 40:1136–1141.

30. Wiktor V, Jonkers HM. Quantification of crack-healing in novel bacteria-based self-healing concrete. Cement & Concrete Composites 33:763–770.

31. Li P, Qu W. 2010. Remediation of concrete cracks by bacterially-induced calcium carbonate deposition. China Civil Engineering Journal.

32. Xu J, Du Y, Jiang Z, She A. 2015. Effects of Calcium Source on Biochemical Properties of Microbial CaCO3 Precipitation. Front Microbiol 6:1366.

33. Zhang Y, Guo HX, Cheng XH. 2014. Influences of calcium sources on microbially induced carbonate precipitation in porous media. Materials Research Innovations 18:S2-79-S2-84.

34. Tittelboom KV, Belie ND, Muynck WD, Verstraete W. 2010. Use of bacteria to repair cracks in concrete. Cement & Concrete Research 40:157–166.

35. Verma SK, Bhadauria SS, Akhtar S. 2013. Evaluating effect of chloride attack and concrete cover on the probability of corrosion. Frontiers of Structural & Civil Engineering 7:379–390.

36. Kügler R, Bouloussa O, Rondelez F. 2005. Evidence of a charge-density threshold for optimum efficiency of biocidal cationic surfaces. Microbiology 151:1341–1348.

37. Yu Q, Zhang R, Deng S, Huang J, Yu G. 2009. Sorption of perfluorooctane sulfonate and perfluorooctanoate on activated carbons and resin: Kinetic and isotherm study. Water Research 43:0–1158.

38. Fan L, Zhang Y, Luo C, Lu F, Qiu H, Sun M. 2012. Synthesis and characterization of magnetic β-cyclodextrin–chitosan nanoparticles as nano-adsorbents for removal of methyl blue. International Journal of Biological Macromolecules 50:0–450.

39. Chiou MS, Li HY. 2003. Adsorption behavior of reactive dye in aqueous solution on chemical cross-linked chitosan beads. Chemosphere 50:0–1105.

40. Dogan M, Ozdemir Y, Alkan M. 2007. Adsorption kinetics and mechanism of cationic methyl violet and methylene blue dyes onto sepiolite. Dyes and Pigments 75:701–713.

41. Zhuang G, Zhang Z, Fu M, Ye X, Liao L. 2015. Comparative study on the use of cationic-nonionic-organo-montmorillonite in oil-based drilling fluids. Applied Clay Science 116:257–262.

42. Takahashi C, Shirai T, Fuji M. 2012. Study on intercalation of ionic liquid into montmorillonite and its property evaluation. Materials Chemistry and Physics 135:681–686.

43. Zerda KS, Gerba CP, Hou KC, Goyal SM. 1985. Adsorption of viruses to charge-modified silica. Appl Environ Microbiol 49:91–95.

44. Chattopadhyay S, Puls RW. 1999. Adsorption of Bacteriophages on Clay Minerals. Environmental Science & Technology 33:3609–3614.

45. Xue W, He H, Zhu J, Peng Y. 2007. FTIR investigation of CTAB-Al-montmorillonite complexes. Spectrochimica Acta Part A Molecular & Biomolecular Spectroscopy 67:1030–1036.

46. Xu W, Johnston CT, Parker P, Agnew SF. 2000. Infrared study of water sorption on Na-, Li-, Ca-, and Mg-exchanged (SWy-1 and SAz-1) montmorillonite. Clays & Clay Minerals 48:120–131.

47. Ross PD, Subramanian S. 1981. Thermodynamics of protein association reactions: forces contributing to stability. Biochemistry 20:3096–3102.

